# MaGuS: a tool for map-guided scaffolding and quality assessment of genome assemblies

**DOI:** 10.1101/032045

**Authors:** Mohammed-Amin Madoui, Carole Dossat, Léo d’Agata, Jan van Oeveren, Edwin van der Vossen, Jean-Marc Aury

**Author notes:** Email addresses: MAM CD LA EVDV JVO JMA.

## Abstract

**Background:** Scaffolding is a crucial step in the genome assembly process. Current methods based on large fragment paired-end reads or long reads allow an increase in continuity but often lack consistency in repetitive regions, resulting in fragmented assemblies. Here, we describe a novel tool to link assemblies to a genome map to aid complex genome reconstruction by detecting assembly errors and allowing scaffold ordering and anchoring.

**Results:** We present MaGuS (map-guided scaffolding), a modular tool that uses a draft genome assembly, a genome map, and high-throughput paired-end sequencing data to estimate the quality and to enhance the continuity of an assembly. We generated several assemblies of the Arabidopsis genome using different scaffolding programs and applied MaGuS to select the best assembly using quality metrics. Then, we used MaGuS to perform map-guided scaffolding to increase continuity by creating new scaffold links in low-covered and highly repetitive regions where other commonly used scaffolding methods lack consistency.

**Conclusions:** MaGuS is a powerful reference-free evaluator of assembly quality and a map-guided scaffolder that is freely available at https://github.com/institut-de-genomique/MaGuS. Its use can be extended to other high-throughput sequencing data (e.g., long-read data) and also to other map data (e.g., genetic maps) to improve the quality and the continuity of large and complex genome assemblies.

## Background

Technical advances and cost reduction in genome sequencing have allowed the completion of numerous genome sequencing projects based on whole-genome shotgun fragments using high-throughput sequencing data and the assembly of these data. The genome assembly process usually involves four main steps: reads assembly into contiguous sequences (contigs), linking of contigs into larger gap-containing sequences (scaffolds), gap closing to fill gaps generated by the scaffolding, and anchoring onto a genetic map to build the final pseudo-molecules. During the second step, end sequences of large fragments (>1 kb) or long reads are aligned to the contigs and the alignment information is used to link contigs into scaffolds. Several commonly used scaffolding programs have been published in the last decade [1]. The efficiency of the scaffolding depends mainly on the diversity and fragment size of the input reads libraries and on the size and quality of the long reads. Typically, 1 to 20 kb libraries are used consecutively during the scaffolding step, which allows repetitive regions of various sizes to be spanned [2]. However, during the alignment step, the presence of repeated sequences creates multiple assembly solutions, which generally causes ambiguities that scaffolder programs cannot untangle. This is often the case in large and complex genomes where repetitive elements are large and cover a large fraction of the genome [3]. To decrease the number of false links, scaffolder programs require a cutoff for the minimum number of read pairs (or long reads) that validate a contigs junction; as a consequence, low-covered contigs are overlooked for scaffold building.

Access to a genome map is a great advantage in obtaining a high-quality genome assembly [4]. Genome maps can also help in detecting assembly errors by revealing discrepancies between the map and the assembly [5] and can provide independent information for evaluating genome assembly quality. Currently, three different types of genome maps can be produced to drive or improve assemblies: physical maps, optical maps, and genetic maps.

Historically, physical maps have been used for large genome sequencing projects to order clones and perform clone-by-clone sequencing, which reduces the complexity of the assembly by sequencing single or pooled clones [6, 7]. Although, this strategy is time consuming and expensive, it remains the best option for high quality genome sequencing of large and complex (polyploid) genomes such as the wheat genome [8]. Recently, the Whole Genome Profiling (WGP™) approach was developed by Keygene NV (Wageningen, The Netherlands) to create an accurate sequence-based physical map starting from a bacterial artificial chromosome (BAC) library [9]. In the WGP method, pooled BAC DNA is digested by a restriction enzyme and after amplification, Illumina technology is used to obtain sequence tags (typically 50 nucleotide sequences flanking the restriction sites). WGP has been used successfully to build physical maps of several plant genomes such as those of wheat [10] and tobacco [11].

Optical maps were used to assemble the *Amborella* [12] and goat genomes [13]. For *Amborella*, this allowed the reordering and super-scaffolding of the draft assemblies and increased their continuity (N50 increased from 4.9 to 9.3 Mb). More recently, the release of the Irys system from BioNano Genomics provided new opportunities to improve the quality and the continuity of genome assemblies [14].

Genetic maps allow the construction of pseudo-molecules by anchoring the assembly on linkage groups that correspond to the chromosomes [15]. Genetic map construction takes advantage of sequence-based genotyping (SBG) [16], genotyping-by-sequencing, and RAD-seq libraries [17] to obtain ultra-dense genetic linkage maps [18]. However, missing data or genotyping errors cause map inaccuracies [19]. Moreover, the physical distance between markers can be very high in genomic regions where the recombination rate is low, which makes it difficult to anchor or orientate scaffolds located in those regions.

Methods used to anchor whole-genome shotgun (WGS) assemblies on genomes have been investigated using several genetic maps to estimate assembly quality, as implemented in MetaMap [5]. The ability of these methods to produce pseudo-molecules also was tested, as reported in Popseq [20] and Allmaps [21]. Allmaps infers the sizes of gaps using the relation between the local recombination rate and the physical distance between two adjacent genetic markers; however, the estimations can be inconsistent considering the inaccuracy of the recombination rate.

Hybrid strategies, combining WGS and genome map data, are likely to help increase the quality of the assembled genome sequence. With this in mind, we developed MaGuS, a modular program that combines a genome map and WGS data. MaGuS can anchor a draft assembly onto a genome map for two applications: quality assessment of a draft assembly by calculating novel metrics, and improvement of the continuity of a draft assembly based on evidence provided by a genome map and high-throughput screening (HTS) data. Here, we detail the MaGuS pipeline and provide an example of its applications using the Arabidopsis TAIR10 genome assembly.

## Methods

### *Arabidopsis thaliana* genome assembly

One 350-bp paired-end (PE) (ERX372154) and two 5.35-kb mate-pair (MP) (ERX372148, ERX372150) Illumina sequence libraries from *A. thaliana* were downloaded from the European Nucleotide Archive (ENA). A total of 95.3 Gb of data were obtained representing a coverage depth of 562X of PE and 170X of MP reads. Adapters and primers were removed from the reads, and low quality nucleotides were trimmed from both ends (quality values lower than 20). Reads were also trimmed from their second N to the end and reads longer than 30 nucleotides were kept. Reads that mapped onto run quality control sequences (i.e., the PhiX genome that is used in Illumina sequencing as quality control) were removed. To decrease the number of sequencing errors present in the paired-end (PE) reads, we applied Musket v1.1 [22] with a k-mer size of 26 ‘-k 26’. We ran Kmergenie v1.5692 [23] on the PE reads to find the best k-mer size for the contig construction step and obtained an optimal k-mer size of 91 bp. For the SOAPdenovo2 [24] assembly, a *de Bruijn* graph was constructed with parameters ‘-K 91 –R’. We selected contigs that were longer than 500 bp.

We used the PE and MP reads in five different scaffolding programs: SOAPdenovo2, SSPACE [25], SGA [26], BESST [27], and OPERA-LG [28]. For SOAPdenovo2, we ran the *map* command with parameter ‘-k 31’, the *scaf* command with parameter ‘–L 500’, and set the minimum number of links in the configuration file as ‘pair_num_cutoff=5’. For SSPACE, we manually set the bowtie k-mer size ‘-l 31’ and ran the program with parameter ‘-k 5’. For SGA and BESST, we first aligned the MP reads onto the contigs using BWA aln [29] with parameter ‘-l 31’. For SGA, the links file was created using the *sga-bam2de* command with parameters ‘-n 5 -m 500 -- mina 31 –k 31’. The *astat* file was generated setting ‘–m 500’. The *scaf* file and the corresponding FASTA file were both created with parameters ‘–m 500’. For BESST, we chose the optimal k-mer size used for the contig assembly as ‘-K 91’ and ran the program with parameter ‘-e 5’. For each program, we selected the scaffolds that were over 2 kb in length. For OPERA-LG, we set the k-mer size for scaffolding with option ‘kmer=91’. The minimum contig size required for the scaffolding step was fixed as 500 bp with the parameter ‘contig_size_threshold=500’. Finally, the number of links to validate a connection between two contigs was assigned with the parameter ‘cluster_threshold=5’.

The source code of QUAST was modified to avoid, as much as possible, the detection of misassemblies (relocation, translocation, and inversion) that correspond to false positives. Because Nucmer generated numerous spurious alignments lower than 5 kb in highly repetitive regions, the minimum alignment length in both parts of a misassembly was fixed as 5 kb. Moreover, the gap or overlap size threshold length was increased to 5 kb to detect relocations. By default, QUAST reports misassemblies found within a scaffold only if at least 50% of the scaffold is aligned. We modified this parameter to report all misassemblies regardless of the aligned fraction of a scaffold.

#### Map-guided scaffolding of genome using MaGuS

First, the WGP tags were aligned to scaffolds using BWA aln [29] and tags with multiple locations were filtered out of the BAM file [30]. We used the resultant alignments to anchor the scaffolds on the genome map and created links between adjacent scaffolds (Figure 1a). However, scaffolds located within other scaffolds, according to the anchoring information, were not considered. More formally, let a tag *t*(*c, r*) be defined by its BAC contig origin *c* and its rank *r*. Let a scaffold *s*((*t*_1_*,p*_1_),(*t*_2_*,p*_2_)*,…*,(*t_n_,p_n_*)) be defined by the n-uplet of a 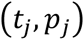 couple, where the tag *t_j_* aligns uniquely at position *p_j_* with *p_j_ ≤ p_j_*_+1_. We define an anchored scaffold 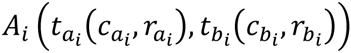 by the origin of the BAC contigs and the ranks of its leftmost and rightmost tags, 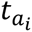 and 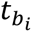, with *r_a_ < r_b_*. We define a map-link between two adjacent scaffolds *a_i_* and *a_j_* only if 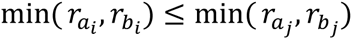 does not include scaffolds located within other scaffolds.

The MP reads were aligned to the assembly using BWA mem [29] and pairs whose mates mapped to different scaffolds were selected. Multiple hits were recorded and mapping possibilities that confirmed a map-link were kept. We estimated the gap size between two adjacent scaffolds from the set of map-anchored scaffolds using the MP fragment size distribution. If the computed gap size was smaller than the maximum expected gap size derived from the MP library, the map-link and the orientation of the two scaffolds were validated. If multiple scaffold orientations were reported by the read mapping, the one supported by the highest number of read pairs was selected. More formally, Let a mapping possibility of a read pair ((*scaf*_1_*,orient*_1_*,pos*_1_),(*scaf*_2_*,orient*_2_*,pos*_2_)) be defined by its scaffold name, orientation, and location of both reads with *scaf*_1_ *≠ scaf*_2_. For each read pair, we calculate the gap size based on the orientation of the two linked scaffolds inferred by each supporting pair, where *len*_1_ and *len*_2_ are the lengths of *scaf*_1_ and *scaf*_2_ respectively, *R* is the read length, and *μ* is the mean of the mate-pair (MP) library fragment size as:

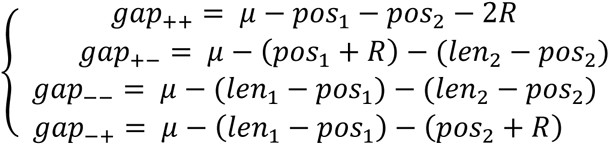

**Figure 1.**
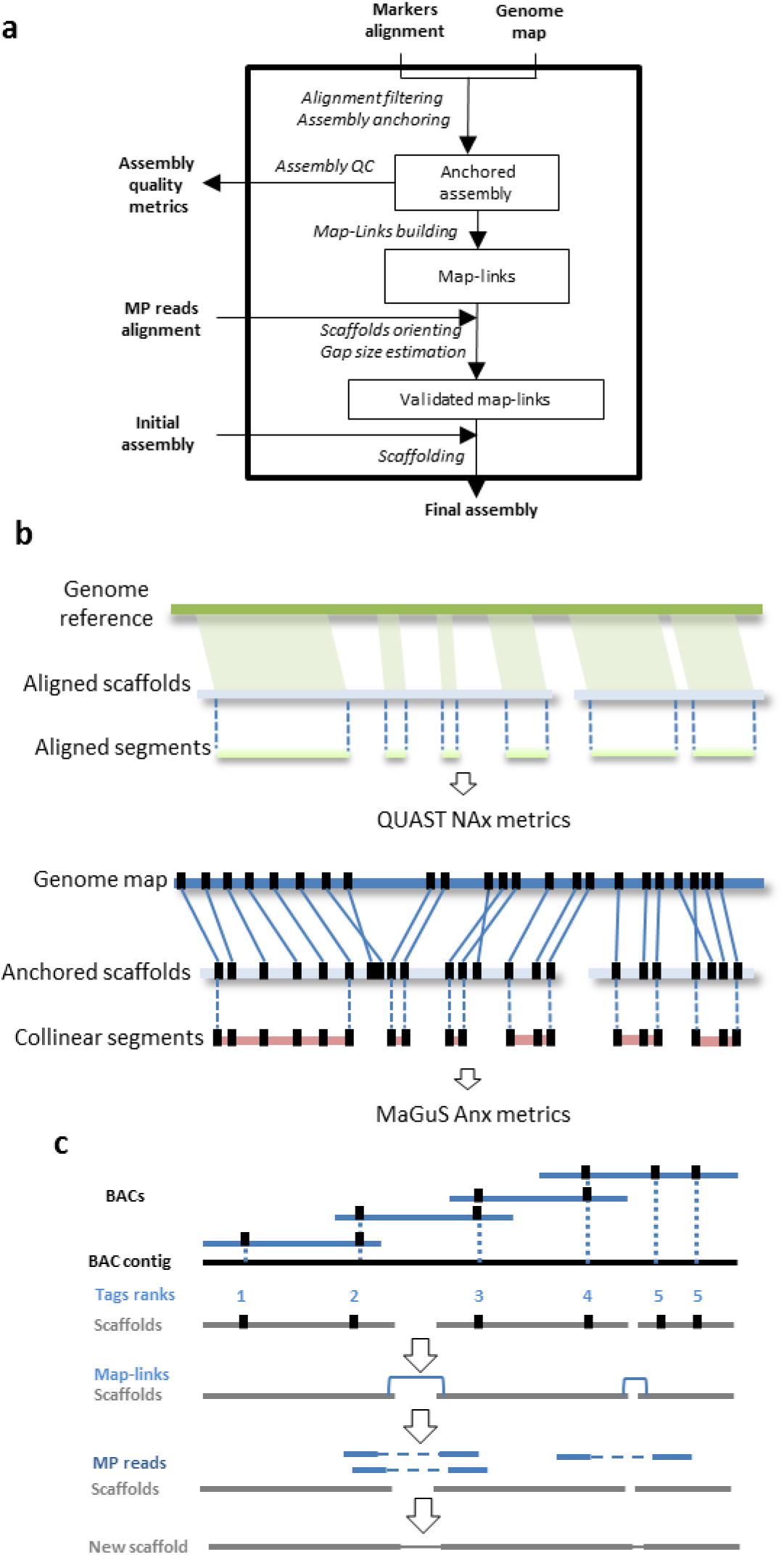
MaGuS pipeline. **a** Flowchart of the MaGuS pipeline. **b** Comparison of the QUAST and MaGuS metrics. **c** Application of MaGuS to WGP data.

We validate the link 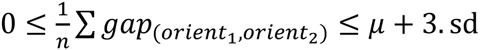, where *μ* and *sd* are the mean size and the standard deviation of the MP library fragment size respectively, and *n* is the number of supporting pairs for the scaffolds link with the following orientation (*orient*_1_*, orient*_2_). Finally, all validated links were formatted for the SGA program to perform the final scaffolding.

#### Analysis of *A. thaliana* WGP data

We used the WGP data produced from the *A. thaliana col-0* BAC library by Keygene (Wageningen, The Netherlands) [9]. WGP tags were ordered by an automated procedure that performed the following steps. First, fingerprinted contig data were read with contig and position information per BAC. Then, BACs were sorted on their left and right positions in the contig and assigned a rank number (identical left and right positions lead to identical ranks). Next, tag information from the WGP tag file was read and occurrences of tags per BAC were listed. For a given contig, a tag position was calculated as the mean value of BAC rank numbers on which the tag occurred. If BAC ranks were too far apart, the tag was identified as an outlier and put aside. The remaining tags were ranked according to their mean BAC rank value, possibly with equal rank scores for equal average BAC rank values.

#### Quality evaluation of genome assembly using MaGuS

We generated new quality assembly metrics from the anchoring based on the commonly used N50 metric (used to evaluate assembly continuity) and the NA50 introduced by the quality assessment tool QUAST (used to evaluate both continuity and quality of assembly using a genome reference [31]). For each scaffold, we defined collinear segments as the fraction of a given scaffold that was correctly organized, i.e., segments anchored with tags that have the same order in the genome map and in the scaffolds (Figure 1b). For a given assembly, the lengths of all these segments were used to calculate the following metrics: An50 (50% of the anchored assembly contains collinear segments with length over An50 bp), AnA50 (50% of the total assembly contains collinear segments with length over AnA50 bp), and AnG50 (50% of the estimated genome size contains collinear anchored segments with length over AnG50 bp). MaGuS also generates Anx, AnAx, and AnGx graphs (based on the Nx graph [2]) that is a plot of the metrics for x values ranging from 1% to 100%.

#### Implementation of MaGuS

MaGuS was implemented in a Perl program based on five modules: *wgp2map*, which performs the anchoring and creates a MaGuS-format map that contains the anchoring information; *map2qc*, which evaluates the quality of the assembly; *map2link*, which creates the map-links between scaffolds; *pairs2links*, which validates the map-links, orients the scaffolds, estimates the gap size, and creates a link.de file; and *links2scaf*, which runs the SGA scaffolding programs and creates the final assembly.

## Results and discussion

### Arabidopsis genome assembly and quality evaluation using MaGuS

PE reads were assembled into contigs with SOAPdenovo2. Then we generated five assemblies using five scaffolding programs (BESST, SSPACE, SOAPdenovo2, SGA, and OPERA-LG) with PE and MP reads. The BESST assembly had the highest continuity (N50 = 1.3 Mb) followed by OPERA-LG (N50=1.27 Mb), SSPACE (0.98 Mb), SOAPdenovo2 (N50=0.82 Mb), and SGA (N50=0.28 Mb). To evaluate the assembly quality, we aligned the scaffolds against the Arabidopsis TAIR10 reference genome with Nucmer [32] using the QUAST pipeline [31] (see Additional file 1 for details). We found that although BESST and OPERA-LG created scaffolds that had longer alignments, they also contained relatively more misassemblies than SOAPdenovo2, SSPACE, and SGA. Based on the QUAST NA50 and NA75 metrics, we ranked the assemblies from the highest to lowest quality as BESST, OPERA-LG, SSPACE, SOAPdenovo2, and SGA.

We used the WGP map to provide a reference-free approach that evaluates the quality of the five assemblies. We applied the *wgp2map* and *map2qc* modules of MaGuS to calculate the length of all collinear segments (Figure1b) and generated Anx values (Table 1, Figure 2a). Considering the MaGuS An50 and the An75 metrics, the ranking of the assemblies was the same as the ranking using the QUAST NA50 and NA75 metrics. The NAx and Anx values were strongly correlated (R^2^ >0.96) for the five assemblies (Figure 2c), which allowed us to consider using the MaGuS Anx metrics to compare assembly quality.

**Figure 2.**
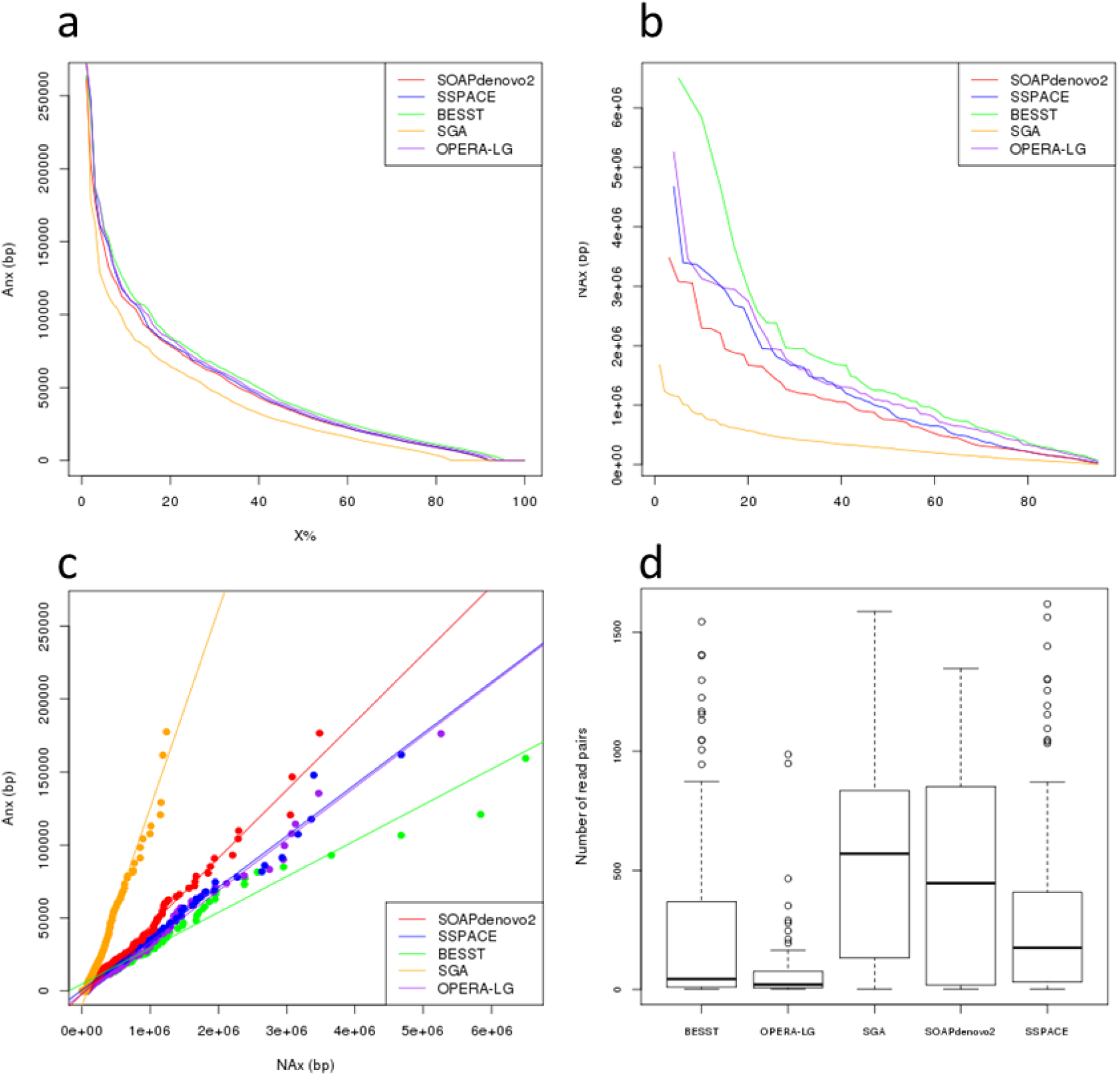
Comparison of MaGuS and QUAST quality metrics for the five assemblies. **a** MaGuS Anx plot. **b** QUAST NAx plot. **c** Correlation between Anx and NAx values.

Figure 3 - Distribution of the number of mate-pairs that validates map-links for the five assemblies

**Table 1.** QUAST and MaGuS quality metrics for the five assemblies

| Assembly metrics | SOAP | SSPACE | SGA | BESST | OPERA_LG |
| --- | --- | --- | --- | --- | --- |
| Assembly size (bp) | 115 319 220 | 116 017 208 | 114 956 386 | 114 996 281 | 116 406 702 |
| N50 (bp) | 821 817 | 982 887 | 284 070 | **1 299 606** | 1 272 891 |
| L50 | 39 | 31 | 115 | **22** | 26 |
| N75 (bp) | 306 051 | 340 070 | 118 727 | **643 037** | 566 836 |
| L75 | 96 | 81 | 270 | **54** | 60 |
| **QUASt metrics** |  |  |  |  |  |
| Number of N's per 100 kbp | 3851.60 | 3000.11 | 4251.16 | 2845.19 | 3139.94 |
| Misassemblies | 9 | 9 | **3** | 23 | 51 |
| Largest alignment (bp) | 3 482 036 | 4 678 885 | 1 680 656 | **6 501 653** | 5 259 610 |
| NA50 (bp) | 757 250 | 926 429 | 276 557 | **1 210 586** | 945 419 |
| NA75 (bp) | 268 694 | 291 099 | 100 235 | **516 026** | 351 844 |
| **MaGuS metrics** |  |  |  |  |  |
| An50 (bp) | 31 217 | 32 028 | 23 164 | **35 466** | 33 908 |
| An75 (bp) | 11 887 | 12 052 | 6 981 | **14 315** | 13 113 |
| R<sup>2</sup> | 0.99 | 0.98 | 0.96 | 0.99 | 0.96 |

The R^2^ values indicate the Pearson correlation coefficients between the QUAST NAx and MaGuS Anx values.

Selecting the appropriate bioinformatics tools to perform genome *de novo* assembly is difficult and often depends on the genome complexity and on the sequencing technology used. The absence of a reference sequence leads automatically to the selection of the assembly that has the highest continuity with no regards to the quality. In the present case, access to a genome map and its use with MaGuS allowed the BESST assembly to be selected as being the most continuous and also the most collinear to the WGP map.

### Arabidopsis genome map-guided scaffolding using MaGuS

We used the five assemblies produced previously to perform map-guided scaffolding through the MaGuS pipeline (Figure 1c). For each assembly, we first created the map-links (i.e., the links between two adjacent anchored scaffolds) and aligned the MP reads onto the scaffolds to validate the map-links by first determining the scaffolds orientation (if the scaffold was anchored by only one tag) and then by estimating the new gaps size (see Methods). The validated map-links were used to build the final scaffolds (Table 2). Only a fraction of the map-links (21.2% to 49.9%) was validated by the MP reads. This limitation was clearly due to the MP library size, and a higher fraction of map-links would certainly be validated using larger MP libraries. Although only a fraction of the map-links were used for the scaffolding, the resulting assemblies showed increases in the N50 metrics ranging from 1.13 to 2.24 times higher and increases in N75 from 1.23 to 2.43 times higher (Table 2). To evaluate the accuracy of this scaffolding approach, we aligned the five assemblies generated by MaGuS onto the Arabidopsis TAIR10 reference genome using QUAST (see Additional file 1). MaGuS generated 86% to 97% correct links for the five assemblies and only a limited number of misassemblies (Table2). The quality of the scaffolds also was confirmed by elevated NA50 and NA75 values. The number of read pairs that validated a map-link had a very wide distribution, from 1 to over 1 000 read pairs (Figure 2c), which showed that MaGuS enabled the scaffolding of both low covered and highly covered regions that corresponded to repetitive regions.

**Table 2.** Assembly metrics after MaGuS scaffolding for the five assemblies

|  | **SOAP** | **SSPACE** | **SGA** | **BESST** | **OPERA-LG** |
| --- | --- | --- | --- | --- | --- |
| Assembly size (bp) | 115 563 956 | 116 414 299 | 115 703 449 | 115 174 685 | 116 556 828 |
| N50 (bp) | 1 350 715 | 1 680 424 | 635 106 | 1 751 177 | 1 442 963 |
| N50 fold change | 1.64 | 1.74 | 2.24 | 1.35 | 1.13 |
| L50 | 23 | 18 | 47 | 18 | 22 |
| N75 (bp) | 509 384 | 646 442 | 288 240 | 787 050 | 695 198 |
| N75 fold change | 1.66 | 1.9 | 2.43 | 1.22 | 1.23 |
| L75 | 58 | 48 | 110 | 42 | 51 |
| Number of N's per 100 kbp | 4 055.34 | 3 331.14 | 4 869.38 | 2 995.68 | 3 264.70 |
| Largest alignment | 5 012 555 | 7 708 756 | 3 361 051 | 6 902 343 | 5 597 743 |
| NA50 | 1 187 620 | 1 455 792 | 579 394 | 1 407 579 | 1 258 868 |
| NA50 fold change | 1.57 | 1.57 | 2.1 | 1.16 | 1.18 |
| NA75 | 354 088 | 508 625 | 215 751 | 609 320 | 560 902 |
| NA75 fold change | 1.32 | 1.75 | 2.15 | 1.18 | 1.59 |
| Total misassemblies | 23 | 19 | 19 | 36 | 62 |
| Magus misassemblies | 14 | 10 | 16 | 13 | 5 |
| Number of map-links | 534 | 481 | 1 034 | 371 | 368 |
| Number of MP-validated links | 209 (39.14%) | 214 (44.49%) | 516 (49.9%) | 93 (25.07%) | 78 (21.2%) |
| Number of correct MP-validated links | 195 (36.51%) | 204 (42.41%) | 500 (48.53%) | 80 (21.56%) | 73 (19.83%) |
| False positive rate | 6.7 | 4.7 | 3.1 | 14 | 6.4 |

## Conclusions

The method presented here and implemented in MaGuS enabled the evaluation of the quality and the scaffolding of a draft genome assembly using a physical map and HTS data. Its application to Arabidopsis with a WGP map provides a first example of its efficiency in reconstructing a eukaryotic genome. Evaluating the quality of a genome assembly is necessary in order to increase the accuracy of downstream analyses, such as genome annotation or comparative genomic analyses. *De novo* assembly projects often lack a genome reference and different ways to assess the assembly quality have been investigated [2, 33] using either the HTS data used for the assembly or a genome map. The latter remains a very good independent source of information for this task. From this perspective, we developed the *map2qc* module of MaGuS to provide assembly quality metrics. Its application to five Arabidopsis genome assemblies showed that the new quality metrics based on the correctly anchored segments of the assembly gave the same assembly ranking as if a reference genome was available. Existing scaffolder tools encounter issues when dealing with repeat-rich regions. The use of a map overcomes this problem if a contig or scaffold can be anchored onto the map. For large genomes, the sequencing depth of an MP library may result in low covered regions. Users of scaffolding programs often set a minimum cut-off for read pairs required to validate a link between contigs, to avoid assembly errors. The use of a map to guide the assembly allows this cut-off to be lowered without loss of accuracy.

## Availability of supporting data

Arabidopsis Illumina reads can be downloaded from the European Nucleotide Archive (ENA) with the following IDs: ERX372154, ERX372148, ERX372150. The WGP data and MaGuS can be accessed through GitHub at https://github.com/institut-de-genomique/MaGuS.

## Competing interests

The SBG and WGP™ technologies are protected by patents and patent applications owned by Keygene NV (Wageningen, The Netherlands). WGP™ is a trademark of Keygene NV.

## Authors’ contributions

MAM designed the method. MAM and CD implemented the method. MAM, LA, CD, and JVO performed the bioinformatics analyses. EVDV and JVO provided the WGP data. MAM and JMA wrote the manuscript. All authors read and approved the final manuscript.

## Acknowledgments

This work was supported by Genoscope (Évry, France), the Commissariat à l’Energie Atomique et aux Energies Alternatives (CEA), France Génomique (ANR-10-INBS-09-08), and KeyGene NV.

## Additional files

Additional file 1 – supplementary methods

